# Targeted nanopore sequencing by real-time mapping of raw electrical signal with UNCALLED

**DOI:** 10.1101/2020.02.03.931923

**Authors:** Sam Kovaka, Yunfan Fan, Bohan Ni, Winston Timp, Michael C. Schatz

## Abstract

ReadUntil sequencing allows nanopore devices to selectively eject individual reads from the pore in real-time. This could enable purely computational targeted sequencing, however most mapping methods require basecalling, which is computationally intensive. Here we present UNCALLED (github.com/skovaka/UNCALLED), an open-source mapper that rapidly matches streaming nanopore current signals to a reference sequence. UNCALLED probabilistically considers k-mers that the signal could represent, and then prunes the candidates based on the reference encoded within an FM-index. We used UNCALLED to deplete sequencing of known bacterial genomes within a metagenomics community, enriching the remaining species by 4.46 fold. UNCALLED also enriched 148 human genes associated with hereditary cancers to 29.6x coverage using one MinION flowcell, enabling accurate detection of SNPs, indels, structural variants (SVs), and methylation in these genes. Twice as many SVs were detected compared to 50x coverage Illumina sequencing, all verified by whole-genome nanopore and PacBio HiFi sequencing.

## Introduction

High-throughput long-read sequencers from Oxford Nanopore Technologies (ONT) produce millions of reads that are several thousand nucleotides in length in a single 48 to 72 hour run. These reads are able to span regions that are otherwise difficult to resolve using conventional short-read sequencing, offering the ability to produce highly contiguous genome assemblies, even spanning centromeric repeats^1^, identify structural variants with significantly higher accuracy, and sequence tens to hundreds of thousands of full-length RNA transcripts in a single run^2^. Nanopore reads can also be used to identify nucleotide modifications, such as methylation, without any additional library preparation considerations^3^.

Nanopore sequencing operates by measuring ionic current as a nucleotide strand passes through a pore. The specific nucleotides in the pore modulate the current in characteristic ways, which can be used to infer individual nucleotides via basecalling of the raw current signal data. For the R9.4 pore, the current is primarily affected by six nucleotides in the central constriction of the pore, which produce signals ranging from 60 to 120 picoamps (pA). Single molecule current readings at these levels are noisy, making it difficult to determine the identity of an individual k-mer. However, by combining the signal information across multiple overlapping k-mers, state-of-the-art basecallers, such as Guppy, can achieve read identities averaging approximately 90%^4^. However, this process is computationally intensive and requires several days to basecall on a high-end multicore CPU. A high-yield run can take well over 24 hours to complete even with a GPU (graphics processing unit).

The ONT MinION is a hand-held low-cost sequencer which typically produces 10-20 Gbp of data from a single standard flowcell. The low price and portability of the MinION has enabled rapid sequencing in remote areas without the need to ship DNA to a sequencing facility^5^. Reads from the sequencer can be output, basecalled, and analyzed as soon as a run begins, which along with the relatively simple library preparation could make rapid sequencing-based diagnostics widely available^6^. The recently released Flongle (flowcell dongle) further improves the accessibility of nanopore sequencing by enabling the use of less expensive flowcells, although this reduces the MinION’s throughput to ~2Gbp. Though this throughput has enjoyed a steady improvement since the initial release of the instrument in 2014, many applications require higher depth, making targeted sequencing necessary. Notably, 20Gbps of data is only approximately 6.6x coverage of a human genome, which is insufficient for most forms of variant calling thereby increasing costs for whole human genome analysis^7,8^.

Typical targeted sequencing methods such as PCR are not suitable for many nanopore sequencing applications. PCR has difficulty amplifying DNA fragments larger than 5Kbp to 30Kbp, which limits nanopore runs that could otherwise produce reads well over 100Kbp. Furthermore, amplification erases nucleotide modifications, which nanopore sequencing could otherwise identify^3,9^. Enrichment methods specifically designed for nanopore sequencing like hybrid capture or CRISPR/Cas9 enrichment alleviate some of these issues, however, they require specialized reagents and extra preparation time^10^. These approaches are also limited in the maximum number and size of regions that can be simultaneously targeted without excessive numbers of reactions and can yield inconsistent amounts of coverage when tiling large regions.

As an alternative, ONT devices have a unique method for real-time targeted sequencing known as ReadUntil, where an individual pore can selectively eject a read while sequencing^11^. This is accomplished by reversing the polarity of the voltage across the specified pore for a short period of time (~0.1s) to eject the molecule and allow a new sequencing read to begin sooner. If one can identify reads that are not of interest and eject them quickly enough, this can enrich the sequencing for targeted regions via a purely computational technique. ReadUntil is more effective with longer reads because each ejection avoids sequencing more nucleotides than if the reads were shorter. For example, the longest reported Nanopore read exceeded 2Mbp in length, and required over 1 hour of sequencing^12^. If this read originated from an off-target region, the sequencing capacity of that pore is effectively wasted for the entire hour, while ReadUntil could have reclaimed that capacity within a few seconds.

In addition to enriching known targets, ReadUntil can instead be used to deplete sequencing of uninteresting or unwanted regions. For example, this could be used to exclude the sequencing of a known microbe in a metagenomics sample or exclude high copy organelles from a plant or animal sample. This is analogous to CRISPR/Cas9-based methods which deplete unwanted sequences^13^, where again ReadUntil has the benefit of not requiring additional library preparation and can uniformly deplete entire genomes as needed. The dynamic nature of ReadUntil could also be utilized to deplete certain sequences after they have been sequenced to a desired depth, which could be useful in metagenomic applications and genome assembly.

A MinION device can sequence up to 512 molecules at a rate of 450 nucleotides per second, requiring a very fast algorithm to effectively enrich regions of interest with ReadUntil. Previous work to enable ReadUntil sequencing used a signal level analysis technique called dynamic time warping to align raw nanopore signal to an *in silico* signal representation of a reference sequence, but this method does not scale to references larger than tens of kilobases as the runtime is quadratic in the length of the sequence^11^. Others have attempted basecalling followed by mapping with a DNA aligner, but basecalling is computationally expensive^14^, and most basecallers are designed to work with fully sequenced reads and require a sizable amount of input signal to output a sequence. A ReadUntil method should ideally be fully streaming, meaning it can continuously refine its classification as more signal is produced. It would also be desirable to have a method that can continue to scale as yields increase with highly parallel devices like the PromethION.

To address these issues, we have developed UNCALLED, the Utility for Nanopore Current ALignment to Large Expanses of DNA, with the goal of mapping streaming raw signal to DNA references for targeted sequencing using ReadUntil. UNCALLED uses the FM-index^15^ to search for sequences in a DNA reference that are consistent with possible k-mers that the raw signal could represent (**Fig. 1a**). It first converts the raw signal into events, which are stretches of signal that approximate k-mer boundaries, and then calculates the probability that each event matches each possible k-mer using a probabilistic model released by ONT. High-probability k-mers are used as a query in a novel FM-index search algorithm developed for UNCALLED which considers all possible sequences and locations for each event as the mapping progresses (**Supplemental Fig. S1**). The probability cutoff used to decide if a k-mer should be considered is dynamically adjusted depending on how many locations a potential sequence could map to, which maintains both high accuracy and high speed when mapping the noisy signal data (**Supplemental Fig. S2**). Finally, a seed clustering algorithm filters out false positive locations by grouping seeds together that map to consistent positions, and a final mapping is reported once a single location has sufficiently more support than the alternatives. We show that UNCALLED can map reads to collections of whole bacterial genomes as fast as a full MinION flowcell can produce reads, and can enrich target genomes by several fold compared to a control. We also show UNCALLED can enrich sequencing a panel of 148 human genes associated with hereditary cancer to a mean of 29.6x on-target coverage, compared to 5.3x coverage on a matched control flowcell, which enables highly precise and sensitive detection of single nucleotide variants, small insertions and deletions, structural variants (SVs), and DNA methylation. Notably, SV calls from enriched UNCALLED reads have 100% concordance with whole-genome ONT and PacBio HiFi sequencing, and detect more than twice the number of SVs compared to whole-genome Illumina sequencing.

**Figure 1.**
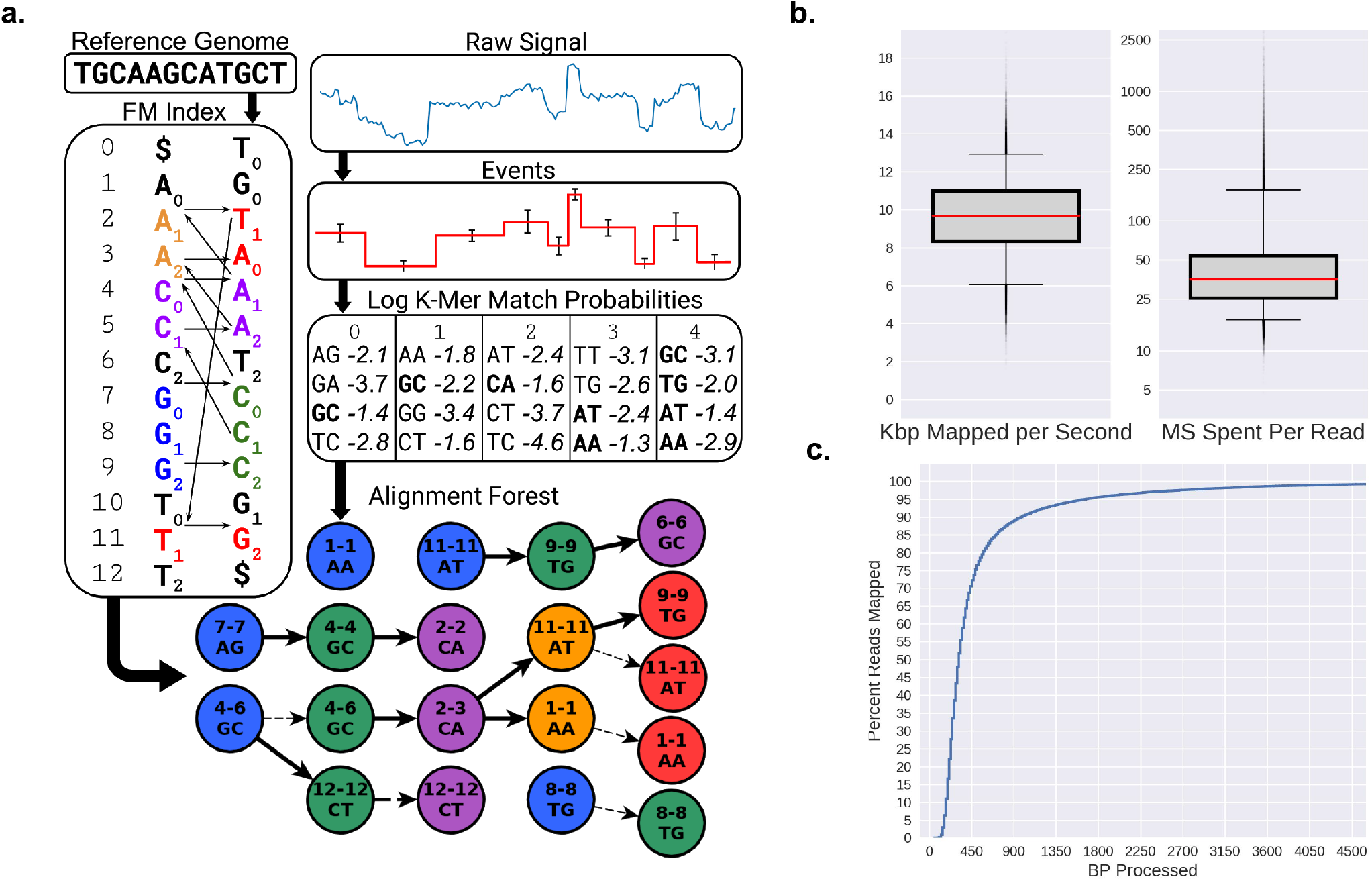
UNCALLED algorithm and performance on *Escherichia coli* data. **(a)** Overview of the algorithm: inputs are an FM index built from the DNA reference, and the raw nanopore signal. The signal is converted to events, and the log probabilities of events matching each k-mer is computed. All paths through the FM-index that are consistent with k-mers that match each event above a threshold are searched, conceptually forming a forest of trees (**Supplemental Fig. S1** for more details). **(b)** Boxplots showing the speed of UNCALLED mapping *E. coli* reads to the *E. coli* K12 reference genome in kilobases per second (left) and total number of milliseconds required to map reads (right). **(c)** Percent of the mapped reads that can be confidently placed within a certain number of basepairs of sequencing. Note the ONT MinION sequences at approximately 450bp per second. Only reads that were mapped by UNCALLED are considered in **b** and **c**.

## Results

### Mapping *Escherichia coli* reads

To measure the accuracy and efficiency of UNCALLED we mapped the raw signal of 100,000 *Escherichia coli* reads to the *E. coli* K12 reference genome using a single 3.0 GHz core. The reads were previously sequenced using a MinION^3^ and have an average length of ~5Kbp. Only the first 30,000 events (~15Kbp worth of signal) at most were considered for each read, which is a default cutoff when using UNCALLED to map previously sequenced reads to avoid spending too much time on exceptionally long reads. Of the reads that mapped, UNCALLED processed a median of 10 kilobases worth of signal per second, the equivalent of 22 actively sequencing pores per thread, and most reads were mapped in under 50 milliseconds (**Fig. 1b**). Of the reads that UNCALLED successfully mapped, 75% were mapped within one second’s worth of sequencing (450bp), which is the amount of signal that the ReadUntil API provides per chunk (**Fig. 1c**). To estimate accuracy we used minimap2^16^ alignments of the basecalled reads as a ground truth: reads that map to the same location are classified as True Positives (TP), reads that neither tool map are True Negatives (TN), reads that are either not mapped by minimap2 or were mapped to a different location than UNCALLED are False Positives (FP), and reads that UNCALLED did not map but minimap2 did are False Negatives (FN). The overall accuracy (TP+TN) of UNCALLED by this analysis was 93.7%. We found that the quality scores (Q scores) of the false negative and false positive reads are much lower than the true positives (**Supplemental Table S1**). Furthermore, 75% of “false positives” consisted of reads that were not aligned by minimap2, meaning the UNCALLED location could be correct and minimap2 could not find it. In addition, of the false positives that minimap2 did align, 92% are explained by repeats: according to nucmer ^17^ self-genome alignments, the reference location that UNCALLED mapped to was a high-identity repeat of the minimap2 reference location. Without repeat masking this type of error is unavoidable if ReadUntil mapping is the goal, since we attempt to find a position based on as little of the read as possible, while minimap2 can consider the full sequence of the read.

### Mapping a mock microbial community

Next, we tested UNCALLED’s ability to map to a collection of genomes using reads from the ZymoBIOMICS High Molecular Weight DNA Mock Microbial community (“Zymo HMW”) containing DNA from seven bacterial species and one yeast (**Supplemental Table S2**). For this experiment, we analyzed 100,000 reads with an average length of 16Kbp we sequenced using a MinION (Full-Flowcell 1 control data). We mapped the signal data from these reads to a 41Mbp reference containing all genomes using UNCALLED, which mapped approximately six kilobases per second with 94% accuracy compared to minimap2 alignments of the basecalled reads (**Fig. 2a**). To test UNCALLED’s performance on different genomes, we used the minimap2 alignments to determine which reads map to each species, and then mapped each collection of reads to their corresponding reference using UNCALLED (**Supplemental Table S2**). The mean read lengths vary between ~11-21Kbp depending on the species, likely because of extraction bias, which skews the mapping rates since UNCALLED is more likely to find a confident mapping location for longer reads. When considering just reads longer than 5kbp long, UNCALLED performs similarly on all bacterial genomes. Mapping to the *S. cerevisiae* genome is ~24% slower compared to the average bacterial genome due to more repetitive sequence than the other references. However, using two iterations of the k-mer masking method described below restores the mapping speed (**Construction and Masking of a Cancer Gene Panel Reference**).

**Figure 2.**
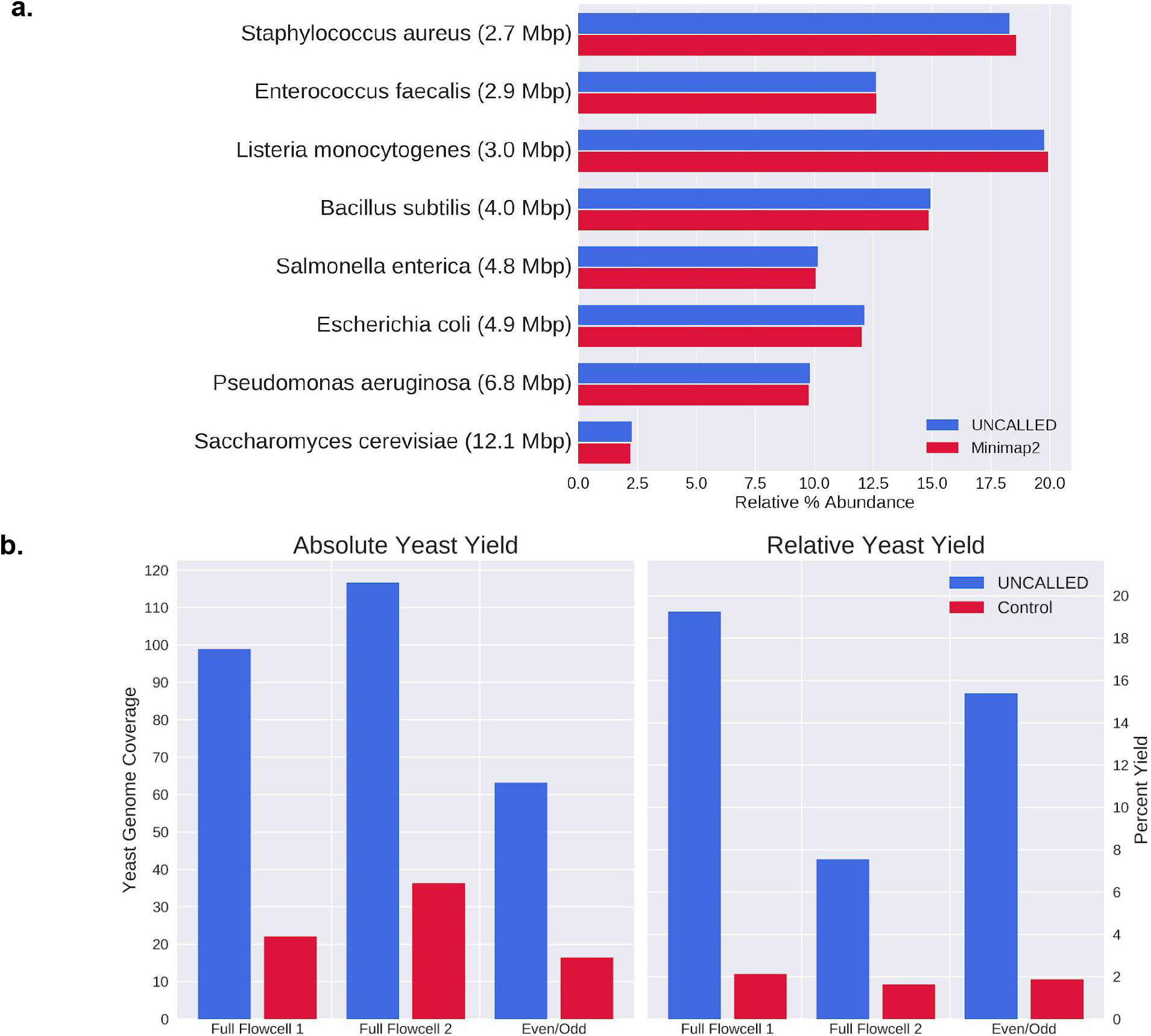
UNCALLED results on the Zymo mock microbial community **(a)** Barchart of the relative abundances of each genome based on mapping reads from a control run to all references using UNCALLED and minimap2. **(b)** Results of UNCALLED ReadUntil depletion of bacterial genomes in order to enrich yeast sequences, including (left) a barplot of the coverage of the yeast genome in the UNCALLED and control runs, and (right) a barplot of the percent of the yield from the yeast genome in the UNCALLED and control runs.

### Bacterial genome depletion

Our first ReadUntil experiment was bacterial genome depletion on the Zymo HMW sample. Here, we used UNCALLED to map signal data to a 29Mbp reference containing all seven bacteria and ejected any reads that mapped within the first ten seconds of signal, with the goal of enriching the yeast sequence which was not included in the reference database. We performed three such runs: two “Full-Flowcell” runs and one “Even/Odd” run. The full-flowcell runs each used two flowcells running in parallel: one sequencing with MinKNOW running in a normal configuration as a control and one with UNCALLED mapping and ejecting reads from all channels. The flowcells were selected to be similar quality based on MinKNOW’s “check flowcell” feature, and the samples were prepared side-by-side and mixed prior to loading. The even/odd run used a single flowcell, where UNCALLED only monitored the even numbered channels and the odd channels were used as a control. This type of control has been used in previous ReadUntil applications^11,14^. Once each run finished, all reads were basecalled with guppy and mapped to a reference containing all Zymo genomes with minimap2 in order to classify reads as originating from yeast or bacteria (**Fig. 2b)**. Note that we classified reads that were not mapped by minimap2 as non-yeast reads, which may underestimate the on-target yield. All UNCALLED runs kept over 99% of yeast reads and ejected between 90% and 96% of bacterial reads, 75% of which mapped within the first second. The absolute enrichment of yeast sequence for these experiments ranged from 3.19 to 4.46 fold (**Supplemental Table S3**).

The enrichment of the even/odd run falls between that of the two full-flowcell runs, implying that this control accurately estimates the enrichment of a full flowcell. Many factors contribute to the variability in the enrichment between runs, including read lengths, reduction in ReadUntil yield, and delayed ejections. In particular, the Full-Flowcell 2 control run had approximately half of the average bacteria read length of the control Full-Flowcell 1 run, which was a major source of the enrichment difference between these runs (**Supplemental Table S3**). We also noted that not all ejections occur as soon as the ReadUntil API request is sent, particularly on high-yield runs. In the Full-Flowcell 1 and Even/Odd runs most reads are ejected within one second of the API call, while in the Full-Flowcell 2 run most ejections are delayed by more than four seconds, increasing the amount of off-target DNA sequenced (**Supplemental Table S3**). Ejections are more delayed early in each run when more pores are alive and actively sequencing. This suggests the ejections may be delayed when too API calls are made at the same time, or when the sequencer is generally overloaded when many reads are being sequenced at once.

It is worth noting that although the on-target yield is consistently higher in the UNCALLED runs compared to the controls, the overall UNCALLED yield is reduced to between 44% and 69% of the overall control yield (**Supplemental Table S3**). Some of this reduced yield can be explained by the short period of time that a pore is empty between sequencing two reads. This gap is not significantly longer when a read is ejected compared to when it finishes normally, but the large number of ejections increases the amount of time that each pore is empty. In the Full-Flowcell 1 experiment, the average channel in the control run was empty ~20% of the time, compared to the average UNCALLED channel which was empty ~32% of the time (excluding time after channels produce their final read and between mux changes). These short gaps are unavoidable, but they do not fully explain the reduced ReadUntil yield. Inspection of the minute-by-minute channel activity throughout sequencing (duty time) shows that the number of functional channels reduces faster in ReadUntil runs, however the long-term lifetime of channels is not shorter than the control, suggesting that pores are temporarily becoming inactive (**Supplemental Fig. S3**). One potential explanation is that ejections cause more pore blockages which make pores unable to sequence reads for extended periods of time. This could be caused by single-stranded DNA on the trans side of the pore self-binding and “clogging” pores, or simply because a larger number of reads are sequenced which increases the chances that a pore will be blocked. Such blockages could possibly be cleared with a nuclease flush, which has been shown to improve yield for other ONT human genome sequencing projects (also see below)^18^.

### Construction and Masking of a Cancer Gene Panel Reference

We next tested UNCALLED’s ability to map to a large collection of human genetic loci. For this, we evaluated an 18.6Mbp subset of the human genome containing 148 genes associated with hereditary cancer from the Invitae cancer panels^19^. These panels consist of curated sets of genes with variants known to increase the risk of developing cancer and are widely used for clinical assessment of disease risk. Our 148 gene panel includes all primary and preliminary-evidence genes from every available organ system panel (**Supplemental Table S4**). This reference was built by extracting all exons, introns, and 20Kbp of intergenic flanking sequence upstream and downstream of each gene from GRCh38 (40Kbp of flanking sequence total). The flanking sequence was included so that reads which start outside but could extend into a gene would be mapped, and so that nearby regulatory elements such as promoters could be covered. To estimate the accuracy mapping to this reference, we used minimap2 to map all 15.7 million reads (mean read length=8,484bp) from the nanopore WGS consortium release 6^20^ to GRCh38 and identified reads that substantially overlap with at least 94% identity any of the 148 genes. We then mapped these reads to the 148 gene reference using UNCALLED to estimate the true positive (TP) rate, which was ~81.71%, substantially lower compared to the bacterial references shown thus far (**Supplemental Table S2)**. We also mapped 200,000 random reads from the WGS consortium excluding the TP reads to estimate the false positive (FP) rate, which was ~1.14%.

We hypothesized that the TP rate was reduced in the 148 gene reference was because the human genome contains much more repetitive and low-complexity sequence than bacterial references, meaning UNCALLED must consider many k-mers for certain signals and therefore uses stricter probability thresholds which make it less likely to find matching seeds. In an attempt to alleviate this we masked the most common 10-mers within the 148 gene reference using an iterative process developed for UNCALLED (see **Methods**). Using 30 iterations of this k-mer masking process raised the TP rate to a level greater than any bacterial reference, however it also raised the FP rate higher than any bacterial reference (**Supplemental Table S5**). Close inspection of these FP reads revealed that they originated from sequences in the 148 gene reference that also occur elsewhere in the human genome. We therefore developed a secondary masking procedure which masked exact repeats greater than 50bp long which occur at least five times in the human genome (see **Methods**). This reduced the FP rate to 1.52% and achieved a 91.60% TP rate, comparable to our bacterial results (**Supplemental Table S5**).

### Cancer Gene Panel Enrichment

With the mapping accuracy established, we next used UNCALLED to enrich for these cancer genes during a MinION sequencing run of the widely used GM12878 cell line. During this analysis, we used the 148 gene reference with 30 iterations of k-mer masking and external repeat masking described above. During this run, we ejected all reads that *did not* map to the gene panel within the first three seconds on one flowcell and used a second full-flowcell as control. We hypothesized that nuclease flushes could unblock pores that may be “clogged” after an attempted ejection, which could be a substantial source of reduced UNCALLED yield as previously discussed. So, we performed nuclease flushes on both runs after 24 and 48 hours and ran each for 72 hours total. The first run resulted in an average coverage over all target regions of 4.0x for the control and 14.3x for UNCALLED, for an overall 3.6 fold enrichment (**Fig. 3a**). Like previous runs, these libraries were prepared without shearing, which typically results in longer reads but reduces overall throughput. We next performed the same experiment but with shearing to 30Kbp, which resulted in an average coverage over all target genes of 5.4x for the control and 29.6x for UNCALLED, for an overall enrichment of 5.5 fold (**Fig. 3a).** In the sheared run, the minimum per-base coverage over all targeted genes for UNCALLED is 7x and over 99.9% of bases have at least 10x coverage, while genes in the control run have several regions with zero coverage and 95.1% of bases have less than 10x coverage (**Fig. 3b**). We noted that after a nuclease flush the fraction of active pores increased in both the UNCALLED and control runs, though the UNCALLED run benefitted more substantially from the flush, supporting the theory that ejected DNA causes pore blockages (**Supplemental Fig. S4**).

**Figure 3.**
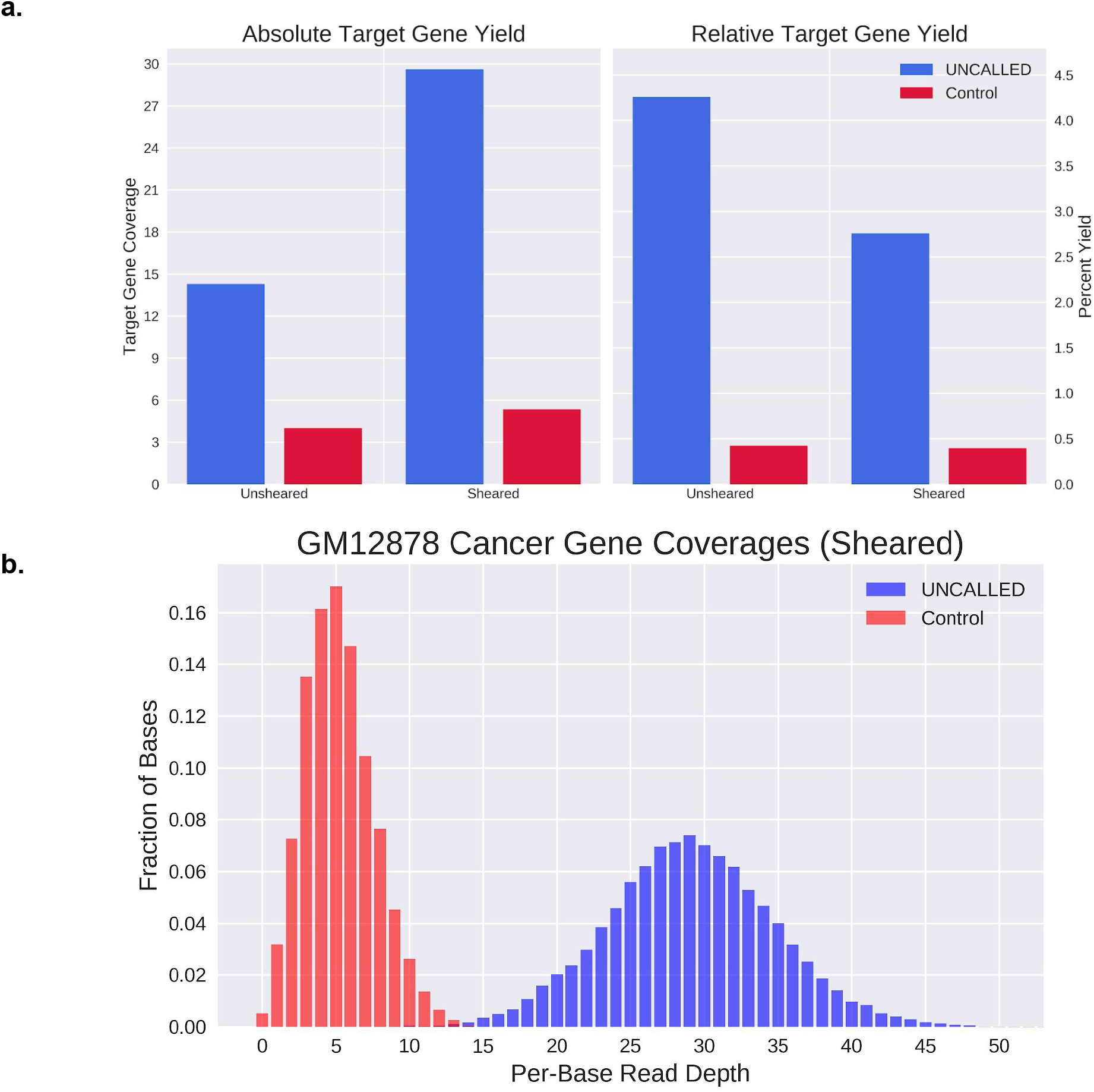
Human cancer gene enrichment using UNCALLED. (**a, left**) Barplot of the coverage over all 148 target genes in the UNCALLED and control runs. **(a, right)**. Barplot of the percent of the yield from the target genes in the UNCALLED and control runs. (**b**) Distribution of per-base coverage over every nucleotide in the target genes in the sheared UNCALLED run. Control ranges from 0x to 15x coverage, while UNCALLED ranges from 7x to 57x coverage.

We next explored applications for the 29.6x coverage sheared UNCALLED reads. We first called single nucleotide polymorphisms (SNPs) and small insertions and deletions (indels) using Clair^21^ on the UNCALLED and control data. For comparison, we also ran Clair on a set of reads with 51.1x coverage over the 148 genes created by combining the two GM12878 control runs, a run sequenced with the same protocol as the GM12878 sheared control, and 37.6x coverage from whole-genome sequencing (WGS) consortium. We compared each call set to the Genome in a Bottle (GIAB) NA12878 small variant truth set^22^ using rtg-tools to compute accuracy metrics^23^. Based on previous work^21,24^, low-complexity regions that are known to substantially reduce small variant calling accuracy were excluded. The precision, recall, and F1 scores of the UNCALLED SNP and indel calls were within one percent of the high-coverage WGS run. In contrast, the control data had less than half the precision, recall, and F1 score (**Table 1a**). The accuracy of indel calls was lower than the accuracy of SNP calls for all datasets, which is consistent with the error profile of ONT reads as shown in previous work^21^.

**Table 1.** Variant calling results over the 148 genes associated with hereditary cancer enriched by UNCALLED. **(a)** Accuracy metrics of single nucleotide polymorphisms (SNPs) and small insertions and deletions (indels) called by Clair. UNCALLED is the 29.6x coverage sheared GM12878 UNCALLED run, and Control is the 5.4x coverage matched control run. ONT WGS is a 51.1x coverage nanopore dataset consisting of WGS consortium reads plus three additional flowcells. **(b)** Structural variants (SV) at least 50bp in length called using the same UNCALLED and ONT WGS reads, plus 30x coverage PacBio HiFi reads and 50x coverage Illumina reads from Genome in a Bottle (GIAB). “Matching UNCALLED” only includes high-confidence SVs detected by each tool. Concordant matching allows an SV to be supported with more sensitive parameters (see **Methods**). (*) Two Illumina high-confidence SVs were inversions, so were not counted among insertions or deletions. However, they overlapped repeats and were not supported by any long-reads, so are likely false positives (**Supplemental Table S6)**.

| a. | Dataset | Count | Precision | Recall | F1 |
| --- | --- | --- | --- | --- | --- |
| SNPs | Control (5x) | 77,346 | 41.9% | 39.4% | 0.406 |
|  | UNCALLED (29.6x) | 12,368 | 92.8% | 97.6% | 0.951 |
|  | ONT WGS (51.1x) | 11,825 | 93.2% | 98.5% | 0.958 |
| Indels | Control (5x) | 25,070 | 37.6% | 23.4% | 0.288 |
|  | UNCALLED (29.6x) | 9,844 | 80.4% | 73.1% | 0.766 |
|  | ONT WGS (51.1x) | 10,374 | 79.9% | 72.7% | 0.761 |

**Table 1.** Variant calling results over the 148 genes associated with hereditary cancer enriched by UNCALLED.
| b. |  | Total SV Count | Insertions |  |  |  | Deletions |  |  |  |
| --- | --- | --- | --- | --- | --- | --- | --- | --- | --- | --- |
|  |  |  | Count | Length (bp) |  |  | Count | Length (bp) |  |  |
|  |  |  |  | Mean | Stdv | Max |  | Mean | Stdv | Min |
| UNCALLED (29.6x) | Confident SVs | 50 | 36 | 196.9 | 175.5 | 974 | 14 | -226.1 | 202.8 | -824 |
| ONT WGS (51.1x) | Confident SVs | 50 | 35 | 206.2 | 178.8 | 964 | 15 | -225.7 | 210.0 | -889 |
|  | Matching UNCALLED | 47 | 34 | 210.1 | 179.9 | 964 | 13 | -241.5 | 220.6 | -889 |
|  | Concordant UNCALLED | 53 | 37 | 197.9 | 177.2 | 964 | 17 | -206.0 | 204.1 | -889 |
| PacBio HiFi (30x) | Confident SVs | 53 | 37 | 199.9 | 176.4 | 992 | 16 | -173.3 | 107.5 | -342 |
|  | Matching UNCALLED | 48 | 34 | 212.4 | 178.8 | 992 | 14 | -181.3 | 110.4 | -342 |
|  | Concordant UNCALLED | 55 | 39 | 203.3 | 175.5 | 992 | 16 | -173.3 | 107.5 | -342 |
| Illumina (50x) | Confident SVs | 25* | 13 | 135.6 | 102.3 | 445 | 10 | -159.5 | 111.3 | -340 |
|  | Matching UNCALLED | 22 | 13 | 135.6 | 102.3 | 445 | 7 | -174.6 | 115.0 | -340 |
|  | Concordant UNCALLED | 25 | 14 | 128.2 | 102.1 | 445 | 9 | -351.2 | 584.7 | -1884 |

With the small variant accuracy established we next called structural variants (SVs) at least 50bp in length in all 148 genes. The 5.4x control run has insufficient coverage for SV calling^7^, so for comparison we detected SVs using the 51.1x coverage ONT WGS run described above, 30x coverage PacBio HiFi reads, and 50x coverage Illumina reads. The PacBio and Illumina datasets were obtained from GIAB. All long-read technologies (UNCALLED, ONT WGS, and PacBio) were called using Sniffles^7^ with minimap2^16^ alignments and the Illumina reads were called using Manta^25^ with BWA^26^ alignments. Strict parameters were used for each dataset to generate sets of high-confidence SVs (**Methods**). These results were compared using SURVIVOR^27^, which showed strong agreement between UNCALLED and ONT WGS SVs (F1=0.94) and between UNCALLED and PacBio SVs (F1=0.93), in contrast to Illumina SVs which matched fewer than half of those predicted by each long-read technology (**Table 1b**). Inspection of high-confidence SVs not matched by SURVIVOR revealed that they all occur in repetitive regions, making it difficult to align short-reads and causing the reported size and location of the SVs vary slightly between the long-reads (**Supplemental Fig. S5 a-e**). Consequently, the apparent disagreements between the long-read call sets were all due to thresholding effects, such as one insertion reported to be 53bp long by PacBio reads (and thus reported as an SV) versus the UNCALLED reads which represented the insertion as 47bp (and thus not reported, **Supplemental Fig. S5a**), while the short-read calls are more fundamentally limited by the challenge of aligning short-reads to repetitive regions. To address the thresholding effects we generated SV calls from each dataset using more sensitive criteria (**Methods**). The SVs in the long-read sensitive calls sets contained matches for every previously unmatched high-confidence long-read SV, demonstrating 100% concordance between all long-read datasets (**Table 1b**).

A total of 56 high-confidence SVs were identified between all long-read technologies (39 insertions, 17 deletions). The sensitive short-read calls resulted in three more concordant SVs compared to strict matching, but still only identified 45% of the SVs detected by long-reads. Four high-confidence Illumina SVs had no support from any long-read technology (two deletions, two inversions), all of which overlap annotated repeats that likely disrupt alignment of the short-reads (**Supplemental Table S6, Supplemental Fig. S5e**).

In order to characterize SVs identified by UNCALLED, we checked for overlap with or similarity to annotated repeats based on RepeatMasker^28^ and simple repeat^29^ annotations from the UCSC genome browser^30^. Insert sequences extracted from PacBio HiFi reads were aligned to the human genome and the primary alignments were checked for overlap with the repeat annotations, which identified repeats in all but one of the 39 insertions (**Supplemental Table S6**). Similarly, all but two of the 17 deletion coordinates overlapped an annotated repeat. Twenty two of the insertions (~56%) and 7 deletions (~41%) were identified as general simple repeats or low-complexity sequences (e.g. “(AT)n” or “G-rich”). Nine insertions (~16%) align to an Alu element, and 5 deletions (~29%) occurred in an Alu element. Interestingly, one of these Alu insertions is located in an exon of the MUTYH gene, an important DNA repair gene associated with colorectal and breast cancers^31^ (**Fig. 4**). The length is consistent with other Alu elements^32^ and it was identified as a heterozygous insertion by all long read technologies but was not detected by the Illumina data. All other SVs occured in intronic regions.

**Figure 4.**
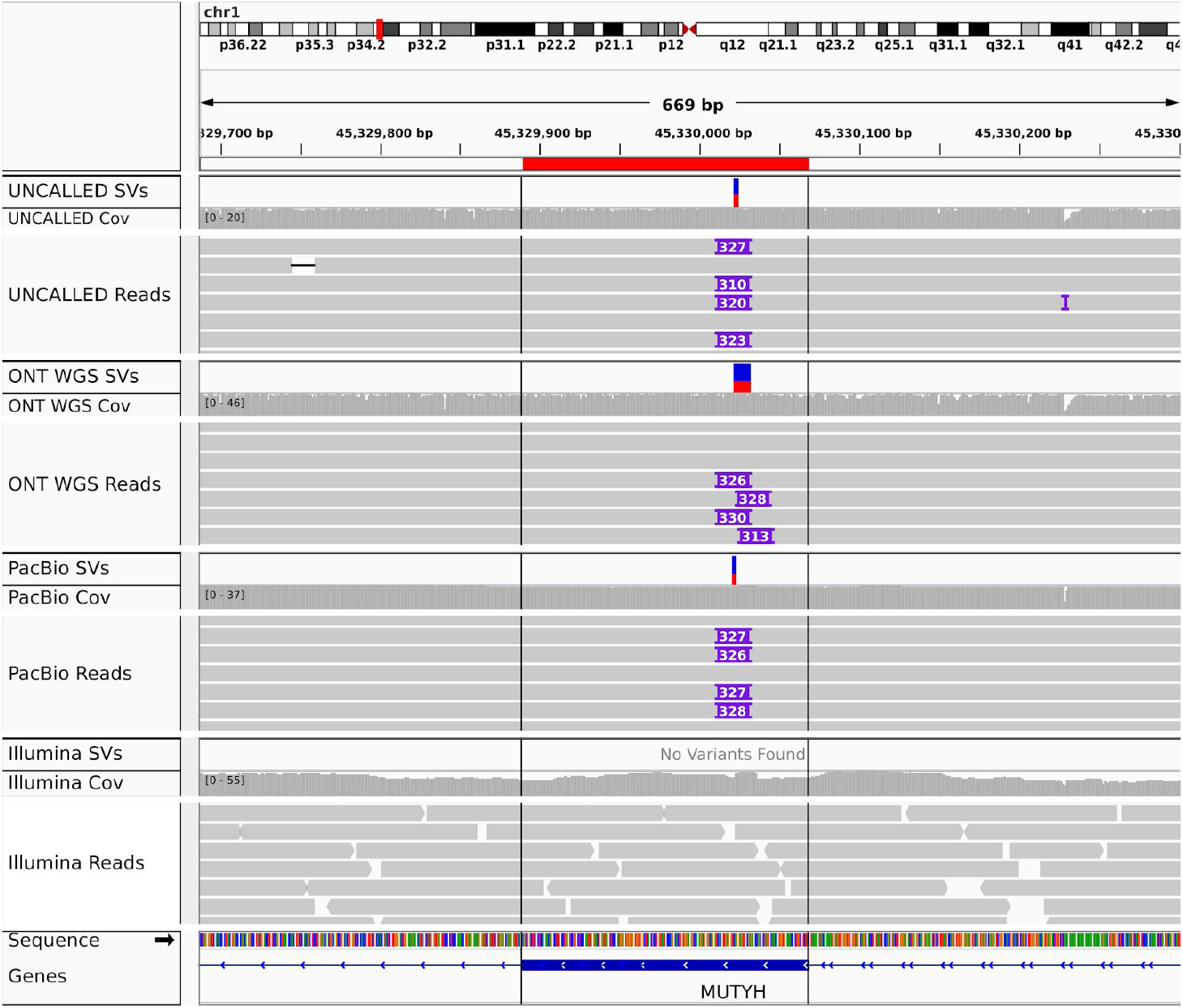
IGV visualization of a heterozygous Alu insertion in an exon of the MUTYH gene detected by UNCALLED, ONT WGS, and PacBio HiFi reads. This was not detected by Illumina reads, likely because short reads cannot span the length of the Alu repeat.

As previously discussed, nanopore sequencing is sensitive to nucleotide modifications, and UNCALLED gives us the depth to characterize them. To demonstrate this we assessed the ability to call methylation over the enriched regions using the UNCALLED sheared run with Nanopolish^3^, again comparing to the 51.1x coverage ONT WGS run described above. Average methylation levels were calculated for 307 annotated promoters with at least 20 CpG sites within the targeted regions for both datasets, resulting in a strong linear correlation (Pearson’s=.96) (**Fig. 5a**). We noted one region near the transcription start site of FANCB, a gene on the X chromosome, where approximately half the reads showed hypermethylation and the other half showed hypomethylation (**Fig. 5b**). This pattern is likely a result of X-inactivation^33^, where the differences correspond to maternal and paternal haplotypes.

**Figure 5.**
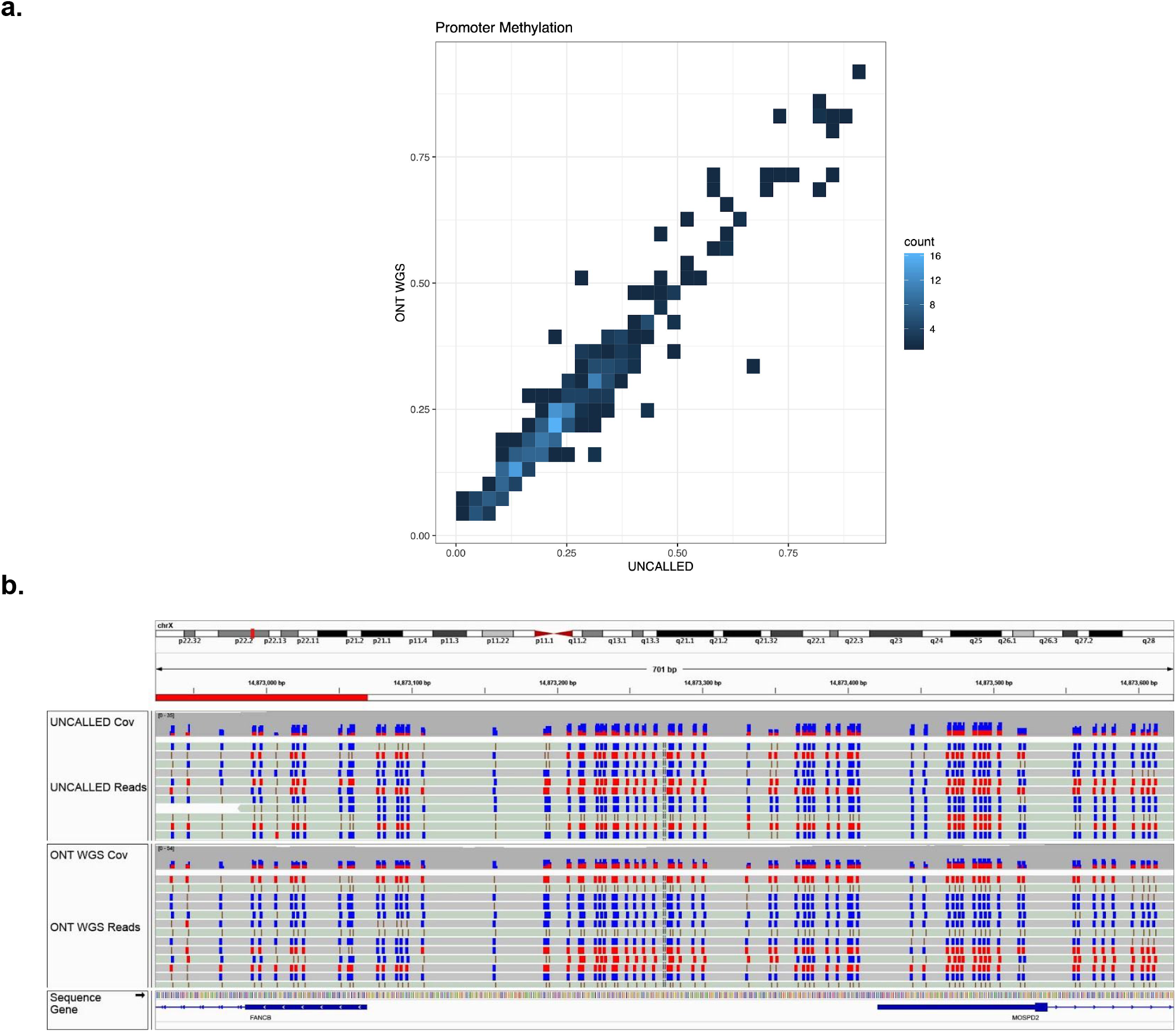
**(a)** Heatmap showing the estimated level of methylation in promoters across the targeted 148 gene regions associated with hereditary cancer in the 29.6x coverage sheared GM12878 UNCALLED run versus a 51.1x whole-genome sequencing (WGS) ONT run. **(b)** IGV visualization at the transcription start site of FANCB on chromosome X. Each individual read tends to be fully methylated or fully non-methylated, likely due to X inactivation.

## Discussion

UNCALLED is a streaming nanopore signal mapper that can accurately map thousands of basepairs worth of signal per second to a reference millions of nucleotides in length. We have demonstrated two ReadUntil approaches: depletion and enrichment. With depletion any reads that confidently map to the reference are ejected. This analysis benefits from UNCALLED’s streaming algorithm, as it does not require a fixed amount of signal to be predefined, and the read can be mapped and hence ejected at any point during sequencing. A use case for depletion, shown by our Zymo community analysis, is to deplete any known bacterial and/or viral contaminants in a sequencing run. Other applications include depleting high copy organelles or plasmids from a sample or depleting the host genome from a host-pathogen sequencing experiments. With enrichment, any reads that *do not* map to the reference are ejected. Here UNCALLED’s ability to map over 90% of reads given less than 3s (~1.3Kbp) worth of signal allows us to accurately make decisions before much of the read has been sequenced. We have demonstrated an important application for this in by enriching all 148 genes used in Invitae hereditary cancer panels to a depth of 29.6x using a single MinION flowcell sequencing GM12878. These data enabled SNP and indel calling more than twice as sensitive and precise as the matched control, and methylation calling that closely matched a 51.1x coverage WGS ONT run. We were also able to accurately identify 56 SVs across all 148 genes with 100% concordance compared to a high-coverage ONT WGS dataset and 30x PacBio HiFi reads, more than twice the number identified with 50x Illumina coverage.

Most SVs detected in GM12878 with the UNCALLED enriched reads were located in intronic regions. While these certainly could have functional effects, exonic mutations are more likely to disrupt gene activity. A single heterozygous insertion was located in an exon of the MUTYH gene (**Fig. 4**), which is a gene involved in DNA repair^31^. Homozygous mutations in this gene are known to cause adenomatous colorectal polyposis, a disease that highly increases the risk of developing colorectal cancer ^34^, and there is some evidence that heterozygous mutations also increase this risk ^35^. Additional analyses and/or functional validation are necessary to determine the effect of this insertion, but a result such as this in a patient would indicate that the individual could be a carrier for adenomatous colorectal polyposis, and could themselves be at a higher risk of developing colorectal cancer.

We have noted several technical issues with the ReadUntil method including read length dependence, pore blockages, and delayed ejections. While UNCALLED is generally more effective with longer reads, longer reads are also associated with lower yield, meaning these factors must be balanced to maximize the on-target yield. This is exemplified in the unsheared and sheared human gene enrichment runs, where the sheared UNCALLED run has a lower percent of on-target yield than the unsheared run, but the absolute on-target yield is higher (**Fig. 3a**). Both ONT yield and read lengths have continually improved historically, so the dependence on long reads is likely to be less of a limitation in the future. Pore blockages are largely eliminated with nuclease flushes, but this requires additional input DNA and preparation time. Blockages could possibly be alleviated by adjusting the voltage applied during ejection, or be avoided by not ejecting reads that map near certain motifs that are likely to self-bind and cause a blockage. ONT has also recently announced plans to incorporate nuclease enzymes directly within the trans side of the pore which could make manual treatments unnecessary^36^. UNCALLED could also eject reads earlier if provided smaller chunks of signal. The ReadUntil API currently only provides signal in one second chunks, while UNCALLED can usually map 75% of reads in less than one second. Reducing this minimum time would allow many reads to be ejected earlier.

UNCALLED is currently limited to mapping to non-repetitive references smaller than ~100Mbp, although the reference can be composed of a collection of many genes and/or many individual genomes. The size limitation is mainly due to the increase of repetitive sequences, since the odds that any sequence appears multiple times in a reference increases with reference size. While we have demonstrated several use cases for UNCALLED, including the enrichment of 148 human genes at once, broadening the types of sequences that UNCALLED can enrich or deplete could be very useful. Optimizations such as further improving the cache-efficiency of the mapping procedure and utilizing SIMD instructions available on modern processors could substantially improve UNCALLED’s performance on any reference. Also, while the repeat masking described here was effective, modifications to the indexing procedure and/or core algorithm could eliminate the need to mask entirely. We also intend to develop a GPU implementation of the UNCALLED algorithm, which could drastically improve the speed, especially to support PromethION sequencing that has more pores available per flowcell and can run multiple flow cells in parallel. Modern ONT basecallers are GPU-based, so we can expect many users to have a GPU, improving UNCALLED’s accessibility. Basecalling reads first, no matter how efficient, levies an additional computational burden requiring more powerful computers, and additional time, meaning ejection is delayed and enrichment lower.

In the future, UNCALLED could be used for additional applications than demonstrated here with little or no modification to the algorithm. For example, UNCALLED could currently enrich or deplete cDNA sequences, which could be useful in avoiding sequencing highly abundant genes or targetting for known gene fusions. ReadUntil with direct RNA sequencing is also possible, although this would require an accurate RNA k-mer model and optimized event detection parameters to account for the different and less stable translocation speed. UNCALLED could also be used in conjunction with other enrichment methods, such as CRISPR/Cas9 enrichment. These methods produce many off-target sequences, which UNCALLED could eject to further improve the amount of on-target DNA. Lastly, UNCALLED can enable many new dynamic applications. For example, an UNCALLED index could be built on-the-fly during a metagenomics sequencing run from the most highly abundant genomes, which could then be depleted for the remainder of the run to increase the coverage for less abundant species. The dynamic nature could also be used to shift the coverage requirements for sequencing below the typical Poisson distribution by selectively ejecting reads from regions of the genome that already have sufficient coverage available. Finally, we also intend to add an optional dynamic time warping (DTW) step to UNCALLED, making it a full-scale signal-to-basepair aligner. This could aid in raw signal applications outside of ReadUntil, such as assembly polishing, identifying nucleotide modifications^3^, and classifying variable number tandem repeats^37^. UNCALLED could improve the sensitivity of such analyses by eliminating the need for basecalling, which can be error prone around such features.

## Acknowledgements

We would like to thank Taher Mun for his contributions on an early prototype of UNCALLED, and Timothy Gilpatrick for providing extracted GM12878 DNA used in the cancer gene enrichment experiments.

This work was funded, in part, by the US National Science Foundation (NSF) grant DBI-1350041 (MCS) and US National Institute of Health (NIH) grant 1R01HG009190 (WT). WT holds two patents currently licensed by Oxford Nanopore Technologies Limited. MCS and WT have received travel funding from Oxford Nanopore Technologies Limited.

## Online Methods

### UNCALLED Algorithm

The core algorithm can be split into three main stages: signal processing, seed mapping, and seed clustering.

***The signal processing stage*** probabilistically decodes the raw signal data into the k-mers they represent. It is based on early Hidden Markov Model (HMM) basecallers, where stretches of similar signal are first collapsed into “events”, which ideally represent the same k-mer^38^. The event detection process can make two types of mistakes, which can be classified as “stays” (multiple events that represent the same k-mer), and “skips” (one event that represents multiple k-mers). Stays are far easier to handle than skips combinatorially: if we know the k-mer associated with one event and we want to predict the next, if we assume the only errors are stays then there are up to five possible next k-mers (extend by A, C, G, or T, or stay), while including skips results in 21 possible extensions (the previous 5 plus extend by AA, AC, AG, AT, CA, etc). We therefore use event detection parameters which are tuned to typically result in ~50% stays and ~1% skips. UNCALLED’s event detector is based on open source event detection code from Scrappie (https://github.com/nanoporetech/scrappie), which uses t-tests over rolling windows to detect when the signal changes significantly to define event boundaries. This code was modified to operate in a streaming manner for UNCALLED. The UNCALLED version produces identical events compared to Scrappie event detection given the same signal and parameters.

Each event is represented by the mean of the signal that it covers. As events are detected, they are normalized so that the mean and standard deviation of a rolling window of events match that of the k-mer model released by ONT ^39^. This is accomplished with a streaming algorithm that computes the mean and variance based on the Welford algorithm^40^ adapted to maintain the rolling window, allowing the normalization to adjust for drift in the signal characteristics throughout sequencing. The default normalization window is 6,000 events long to ensure a robust sampling of all possible k-mers.

After normalization, UNCALLED calculates the probability that each event matches each possible k-mer based on ONT’s k-mer model. This model lists the expected mean and standard deviation of the signal associated with each 6-mer, which is modeled as a normal distribution. To accelerate computational processing, UNCALLED uses a simplified model of 5-mers with little loss of information when computing event/k-mer match probabilities. During signal processing, UNCALLED picks a probability threshold that is dynamically altered depending on how uniquely the seed is mapping, and considers all k-mers which match each event above that threshold (see **Index Probability Thresholds** below).

***The seed mapping stage*** attempts to find relatively short but perfect alignments between the read and the reference genome. UNCALLED uses an FM-index, which is the data structure used by many aligners such as Bowtie ^41^, BWA ^26^, and HISAT ^42^. BWA provides a library for its FM-index, which UNCALLED directly uses to take advantage of its highly optimized construction and querying. The FM-index allows one to find all locations of an arbitrarily long query sequence in a reference, with time essentially constant with respect to the reference size. UNCALLED uses a novel branching algorithm which considers all k-mers that each event can match at each step of the mapping. This algorithm speed is not constant with respect to the reference size due to the branching, but scales much better compared to dynamic time warping.

Given a new read, UNCALLED first finds all locations of all k-mers which match the first event. The FM-index allows an efficient representation of all locations of each unique sequence. For the next event, it checks if that event could match any k-mers “compatible” with any of the k-mers that matched the previous event. A 5-mer is compatible if its first four bases match the previous 5-mer’s last four bases, or if they are the same 5-mer in the case of a stay. Each compatible non-stay k-mer extends the previous sequence by one basepair, and we use the FM-index to find the locations of each extended sequence. This search space conceptually forms a forest of trees, where each possible sequence and the locations of that sequence are represented as a path from a root to a leaf (**Supplemental Fig. S1**). After existing mapping paths are extended, UNCALLED begins new paths by finding the locations of k-mers that match the event but are not represented in the previous paths. Again, the FM-index provides an efficient mechanism to accomplish this. The UNCALLED algorithm proceeds by alternating between extending old paths and creating new ones, and can report a seed mapping when one sequence narrows down to a unique location in the reference.

Storing mapping paths as nodes of trees connected by edges would be computationally intensive due to cache inefficiency when accessing non-contiguous memory. To improve performance, UNCALLED stores each path from a root to a leaf in a “path buffer” (**Supplemental Fig. S6**). Each path buffer stores cumulative log probabilities of each event matching each chosen k-mer, and other information such as the FM-index location and the most recent k-mer matched. When a path branches, the buffer is copied to preserve this information for each extension. The length of the path buffers determines the seed length, and a seed is only reported if the buffer is full and the mean probability over all events in the buffer is above a threshold. When an event is added to a full buffer, the oldest event is erased and all other events shift to make room for the new event, allowing subsequent seeds to build off of previous seeds.

***The seed clustering stage*** separates true alignments from spurious seed matches. Due to the noisy nature of nanopore sequencing, UNCALLED must use very loose thresholds for event/k-mer matches, which produce many false positive seed mappings. We eliminate these false positives under the observation that they will usually map to random locations, while true positives will map to locations consistent with their position on the read. This analysis is complicated by stays, which can occur inconsistently: there are often long stretches of stays followed by many non-stays. We therefore developed a rapid clustering algorithm which groups seeds together if their read and reference coordinates are consistent with each other. Specifically, the distance between read coordinates of adjacent seeds must be larger than the distance between the reference coordinates, but not by more than a factor of 12 by default, which handles most stretches of “stay” events. When a seed is added to an existing cluster of seeds, we update the total number of reference basepairs that the cluster covers, which is used as a measure of support for that reference location. We report a read mapping once the ratio of the best supported location over the next best supported location is above a threshold (default: 1.85 fold), or the ratio of the best supported location over the mean support for all locations is above a different threshold (default: 6.00). The first threshold is sufficient for most non-repetitive regions, while the second threshold is somewhat more repeat tolerant.

### Index Probability Thresholds

UNCALLED uses the BWA library for constructing, storing, and querying the FM index ^26^. After the index is constructed, reference-specific probability thresholds must be precomputed to maintain accuracy and speed for references of different sizes and repeat contents. These thresholds are used to decide which k-mers can be used to extend a path based on the event/k-mer match probabilities (see **the signal processing stage** above). The threshold to be used depends on how many reference locations that the path being extended could currently map to, which is determined by the length of the FM index range associated with the path buffer (FM range size, **Supplemental Fig. S2**). The goal in choosing these thresholds is to limit unnecessary branching in the seed mapping process. A path buffer that could map to many locations (a large FM range size) should use a strict threshold, meaning fewer k-mers are likely to be considered, since each location that a path can map to could form a new branch later in the mapping process (**Supplemental Fig. S1**). As a path gets longer the number of locations to which it could map tends to decrease, and so using more permissive probability thresholds for smaller FM range sizes increases the odds that events extending longer paths will correctly match the k-mers they represent. The correspondence between FM range sizes and probability thresholds must vary depending on the reference, since paths of the same length are likely to have longer FM range sizes as references become larger and more repetitive.

UNCALLED uses a different log probability threshold for every power of two FM range size, and assigns thresholds such that FM ranges with size in the range [2^*s*^, 2^*s*+1^] share the same threshold for any positive integer *s* (**Supplemental Fig. S2b)**. This works well since FM range sizes decrease exponentially as mapping progresses, and it is computationally efficient to index in real-time by computing the log base two. We developed an EM algorithm to assign probability thresholds for intervals of FM index range sizes with the goal of minimizing the total amount of time required to map a read.

Mapping paths are always extended one nucleotide at a time. It is useful to have a correspondence between the number of nucleotides mapped, which we call the “path position”, and the expected FM range size at that position. To find this correspondence we use the FM index to map the reference genome to itself from many random locations (one out of 100 genome locations by default) until a unique location is found, which provides a sampling of the FM range sizes for each path position. We then compute *N*, which is the number of basepairs required to map 95% of positions uniquely (by default), and *M* which is the maximum log FM size for all k-mers in the genome (*k* = 5 for the 5-mer model used by UNCALLED). For every *p* ∈ [*k*, *N*] we compute the mean log_2_ FM range size after *p* basepairs are mapped which we call *F*(*p*), and for every *f* ∈ [1, *M*] we compute the mean of all path positions with FM range size *f*, which we call *P*(*f*). These functions allow us to pick thresholds with respect to path positions and then map those thresholds back to FM range sizes. Assuming we have some function *T* (*p*) which returns a log probability threshold for every path position *p* ∈ [*k*, *N*], we aim to compute a “speed coefficient” *S* which is proportional to the basepairs per second that we expect to map. We can then use an EM algorithm to adjust *T* (*p*) until an optimal *S* is found. *S* is computed based on the mean number of k-mers that we expect to match an event above each threshold, which we call *B*(*t*) (**Supplemental Fig. S2a**, blue line). For each path position, the amount of work UNCALLED must perform is proportional to the number of k-mers we expect to consider at the given threshold multiplied by the number of reference locations that currently considered: *B*(*T*(*p*)) × *F*(*p*).

These values are summed for every path position and normalized to compute *S*:

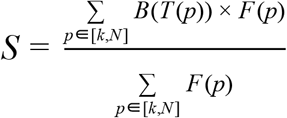

Finally, we must define *T*(*p*) to be some increasing function that has an adjustable variable to optimize with the EM algorithm. Various functions were tested, but the one used in UNCALLED is based on the power function: *y* = *x*^θ^. This function always intersects (0,0) and (1,1) making it easily scalable, and adjusting θ makes it increase faster or slower. This requires some fixed initial and final probability threshold, which we call *t*_0_ and *t*_*N*_ respectively. The default values for *t*_0_ and *t*_*N*_ are −10.00 and −2.25 respectively (both natural log probabilities), which were empirically determined to be the lowest and highest useful thresholds (**Supplemental Fig. S2**). We then define *T*(*p*) as:

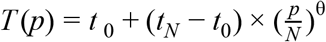

The EM algorithm then works by adjusting θ until *S* reaches the target value. The default target for *S* is 115, which was empirically found to minimize the total amount of time to map reads to various references. Once θ is found we can compute *T*(*P*(*f*)) for all *f* ∈ [1, *M*] to find probability thresholds for every log_2_ FM range size. An example set of probability thresholds computed for the *E. coli* K12 reference genome can be seen in **Supplemental Fig. S2b**.

### Implementation

UNCALLED is available open source at https://github.com/skovaka/UNCALLED. The core algorithm is written in C++, with a Python frontend to interact with the ONT ReadUntil API. UNCALLED uses the BWA ^26^ FM-index to take advantage of its highly optimized construction and querying. It can be run as a standalone read mapper in addition to live ReadUntil, and outputs locations in the PAF (Pairwise mApping Format) introduced by Minimap ^43^. UNCALLED was run on a 24 core 3.0 GHz Intel Xeon Gold 6136. All mapping speeds were measured by running with a single thread. All ReadUntil experiments were run using 48 threads.

### Zymo Bacterial Depletion Experiments

Bacterial reference genomes for the ZymoBIOMICS High Molecular Weight (HMW) DNA Standard were obtained from the ZymoBIOMICS (https://s3.amazonaws.com/zymo-files/BioPool/D6322.refseq.zip). A *S. cerevisiae* draft genome was also included, but this reference was highly fragmented, so the S288C reference genome (NCBI accession GCF_000146045.2) was used instead for mapping with UNCALLED and Minimap2.

The Full-Flowcell 1 UNCALLED run ejected reads throughout the entire sequencing run including during “mux scans” when the sequencer checks pore quality to prioritize pore usage. Subsequent ReadUntil runs did not eject during mux scans in an attempt to improve yield by preventing ejections that might disrupt the MinION’s ability to check pore quality. This did not appear to have a substantial impact on ReadUntil yield, but the feature was left in to not disrupt the mux scans since they account for less than 2% of sequencing time.

### Hereditary Cancer Gene Reference

The 148 genes associated with hereditary cancer were obtained from the Invitae website by extracting the names of all primary and preliminary-evidence genes from every listed panel^19^. The coordinates of these genes in GRcH38 were identified in the Ensembl gene annotation (v98)^44^. The gene coordinates were the extended by 20,000bp on each end to include flanking sequence, and the sequences at those locations were extracted from GRcH38 using bedtools^45^ to obtain an 18.6Mbp reference.

Two forms of masking were performed on the reference containing 148 human genes. The first is an iterative process based on identifying the most common 10-mers that occur within the reference. In each iteration the most common 10-mer is first identified using jellyfish ^46^, and then that k-mer is masked by replacing each occurrence with “N”s. The BWA indexing procedure replaces any “N” with a random basepair, making it highly unlikely that a read will falsely map to ten or more “N”s. In the next iteration the masked reference from the previous iteration is used, meaning the previously most common k-mer will no longer occur, and any k-mers overlapping that k-mer will have a reduced count. This method was also applied to the *S. cerevisiae* genome to demonstrate an improvement in mapping speed.

The 10-mer masking process is effective in increasing the TP rate, but also increases false positives caused by repeats outside of the reference. To identify these external repeats we first extracted all 50bp windows from the 148 gene reference and aligned them to the full human genome using bowtie ^41^, using parameters which find all exact end-to-end matches (−a −v 0). We then found all windows that occur at least five times in the genome and merged them to find all contiguous regions 50bp or larger that occur at least five times. These regions were again masked by replacing them with “N”s.

### Small Variant Calling

SNPs and Indels were called using Clair^21^ with the alternative allele frequency threshold set to 0.2 as recommended for ONT reads. We used the vcfeval command in rtg-tools^23^ to compute precision, recall, and F1 scores. For the truth set we extracted all entries in the Genome in a Bottle (GIAB) NA12878 small variant truth set^22^ which overlap the target genes, and removed variants which overlap low-complexity regions which negatively impact small variant detection benchmarking according to the Global Alliance for Genomics and Health (GA4GH)^21,24^. Variants in these regions were also removed from the Clair outputs. This was done based on regions specified in previous work^21^.

### Methylation Calling

CpG methylation was called using Nanopolish on default settings. Promoter regions, as annotated in the Ensembl regulatory features of GRCh38, within the targeted regions were identified using bedtools^45^. Average methylation frequency was then calculated over these promoters. For read-level visualization, CG positions and methylation calls were annotated in the alignment files (https://github.com/isaclee/nanopore-methylation-utilities).

### Structural Variant Analysis

High-confidence long-read SVs were found using Sniffles ^7^ with a minimum SV length of 50bp and a minimum read support of one quarter of the average coverage: 7 for UNCALLED and PacBio, and 13 for the high-coverage WGS dataset. All other Sniffles parameters were left at their defaults. The minimum read support was chosen to be one quarter of the average gene coverage per sample as used in other studies^7,47^, based on the fact that heterozygous SVs should be represented in approximately half of all reads so that one quarter of reads will capture SVs with high probability ^47^. High-confidence short-read SVs were found using Manta^25^ using the default score cutoffsless. Only SVs that are less than 1Mbp in length, do not include translocations to other chromosomes, and overlap one of the 148 cancer genes were considered for both Sniffles and Manta. The high-confidence SVs were matched using “SURVIVOR merge” with a maximum distance of 1,000bp and requiring the same variant type and strands. Because of the noise in the raw ONT reads, Sniffles split the same SV into two separate SVs in both the UNCALLED and WGS datasets (**Supplemental Fig. S5f**), which SURVIVOR merged in the WGS run but not UNCALLED. Accordingly, this case was manually corrected. Matches for previously unmatched high-confidence SVs were found using more sensitive criteria in Sniffles (minimum length of 30bp, minimum read support of 3, and maximum SV grouping distance of 50bp), and using all Manta SVs regardless of scoring. The maximum matching distance in SURVIVOR was also set to 1,500bp for merging the sensitive call sets.

Deletions were characterized by first intersecting their reference coordinates with all RepeatMasker^28^ entries downloaded from the UCSC Genome Browser^30^ using bedtools^45^. If no overlap was found with RepeatMasker, we searched for overlaps with the “Simple Repeat” track from UCSC Genome Browser, which is based on Tandem Repeat Finder annotations^37^. Insertions were classified by extracting the insert sequence output by Sniffles for the PacBio HiFi SV calls, aligning this sequence to GRcH38 using BWA MEM ^26^, and intersecting the alignments with RepeatMasker and Simple Repeats. No alignment was found for two of the insertions, so these were characterized by their reference coordinates in the same way as deletions.

### Samples

ZymoBIOMICS HMW DNA Standards were purchased from Zymo Research. GM12878 cells were purchased from Coriell Institute and propagated in Dulbecco’s Modified Eagle Media (DMEM) with fetal bovine serum (FBS), penicillin, and streptomycin at 37°C and 5% CO_2_. Cells were then snap frozen in pellets of roughly 2 million cells, and DNA was extracted from the cell pellets using the Nanobind Cells, Blood, Bacteria (CBB) Big DNA Kit from Circulomics according to the manufacturer's specifications. The extracted DNA was then directly sheared without dilution to 30kb using the Diagenode Megaruptor 2. Sheared samples were processed twice in a row on these settings due to the viscosity of the extracted DNA. Before library preparation, short fragments of DNA were depleted from the samples using the Circulomics Short Read Eliminator XS (SRE XS) kit, according to the manufacturer’s specifications.

### Library preparation

All sequencing libraries were prepared using the ONT Ligation Sequencing Kit (SQK-LSK109) without the DNA Control Strand (DCS) or FFPE repair in the end prep step. The initial sample volume was thusly adjusted to 50ul, and Ultra II End Prep Reaction Buffer volume was adjusted to 7ul. Nuclease flush Buffer A was prepared by combining 659ul of ultra pure water, 300 ul of 1M KCl, 30ul of pH 8.0 HEPES buffer, 10ul of 1M MgCl_2_, and 1ul of 2M CaCl. Just prior to loading the flowcells, the libraries for the control run and the UNCALLED run were mixed together, as were the priming buffers for the runs. ONT MinION flowcells (FLO-MIN106) with vR9.4.1 pores were used for all sequencing. Flowcells were selected such that the estimated available pores on the UNCALLED and control runs were within 200 pores of each other (out of ~1,400 to ~1,700 total pores). The runs used in addition to the WGS consortium data for the high-coverage WGS SV calls were the two GM12878 control runs, plus another library sequenced with the same protocol as the unsheared GM12878 control run.

### Data Availability

All sequencing runs are available as an NCBI BioProject under the accession PRJNA604456.

## Supplemental Figures

**Supplemental Figure S1.**
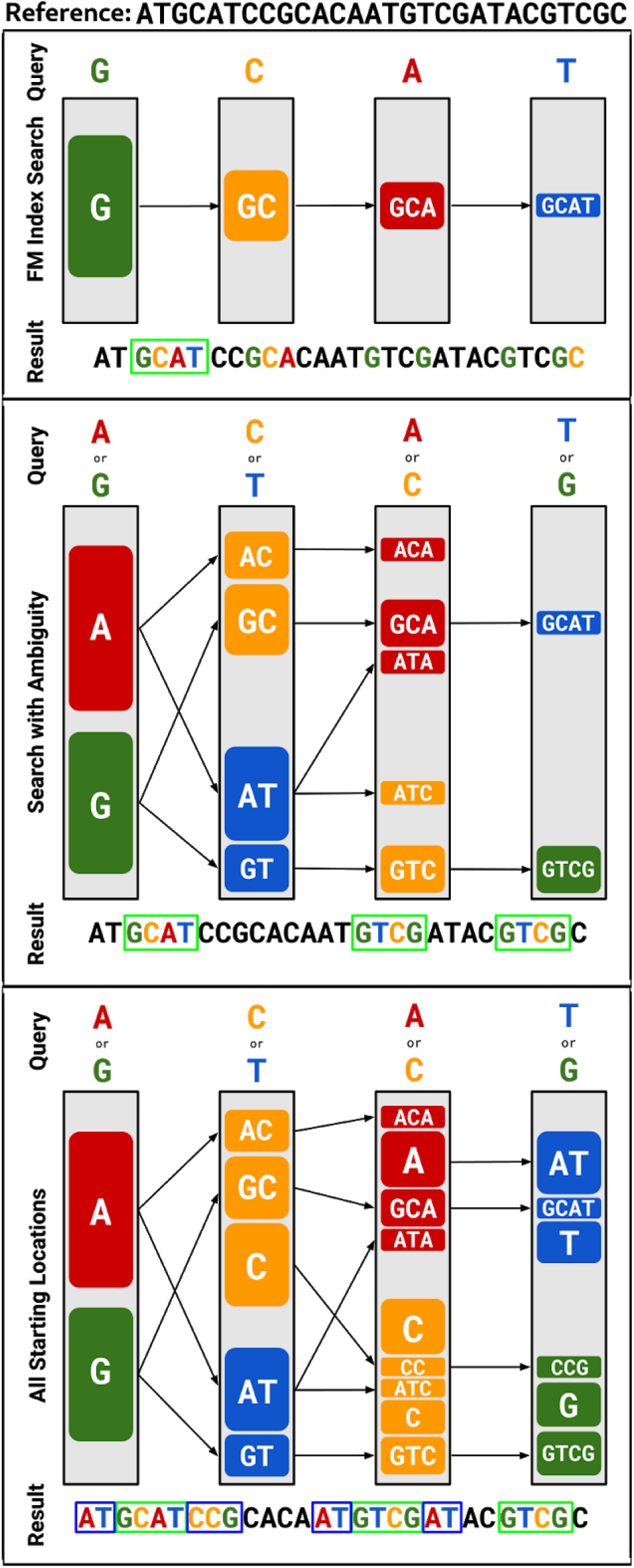
FM Index alignment examples. (**top**) FM index alignment of a standard DNA sequence, where the size of each box represents the number of possible locations. (**middle**) FM alignment of a sequence where every position could be one of two bases. Base ambiguity is analogous to the k-mers we consider for every event. (**bottom**) Same as middle but alignments starting from all positions are found by filling in the gaps between ranges from previous alignments.

**Supplemental Figure S2.**
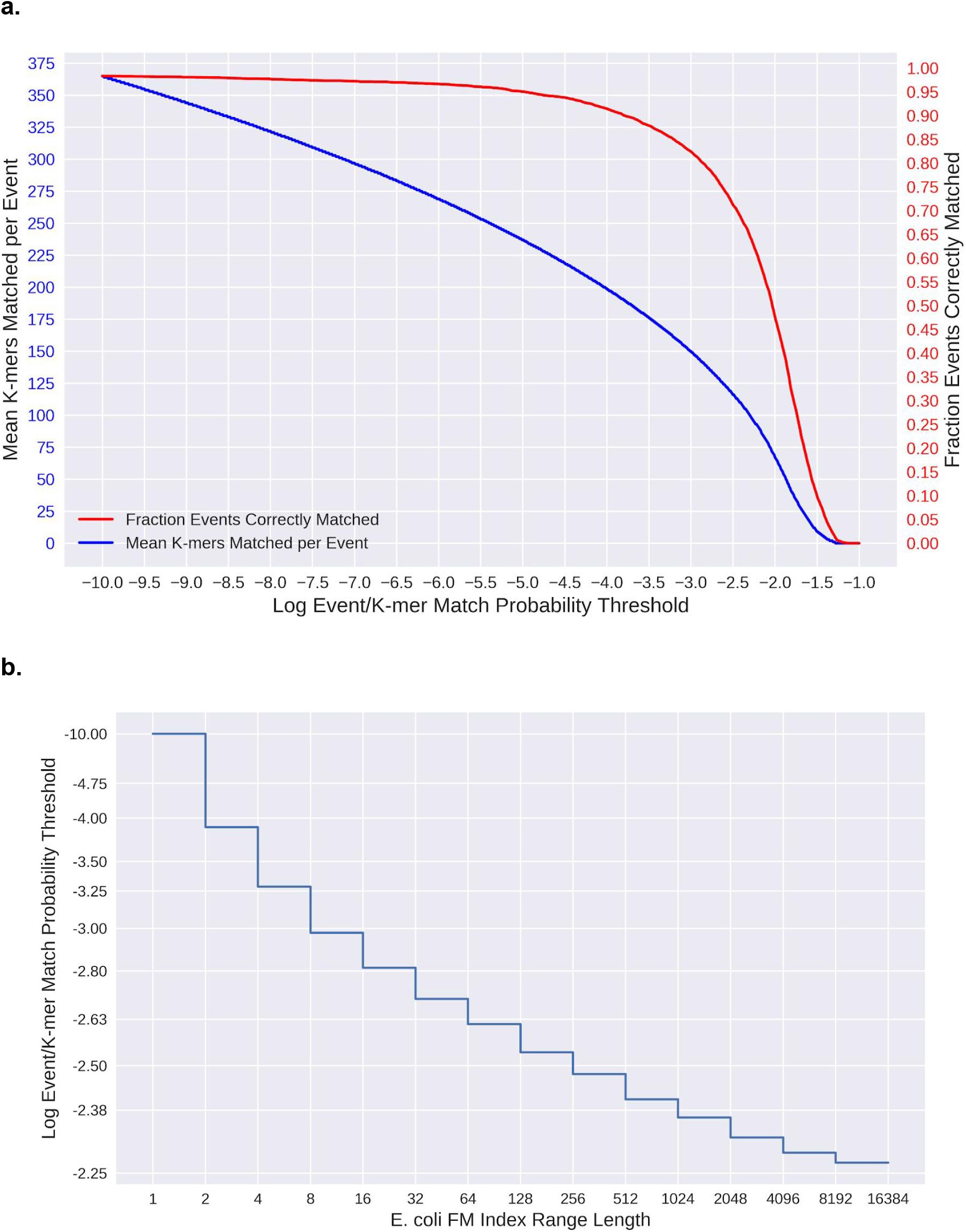
Match Probability Thresholds **(a)** Relationship between natural log probability thresholds (x-axis), the mean number of k-mers that match above each threshold per event (blue), the fraction of events that match their correct k-mer above each threshold (red). The values for r9.4 chemistry are shown here. **(b)** The FM index range lengths assigned to different probability thresholds for the *E. coli* reference. This function varies depending on the reference used.

**Supplemental Figure S3.**
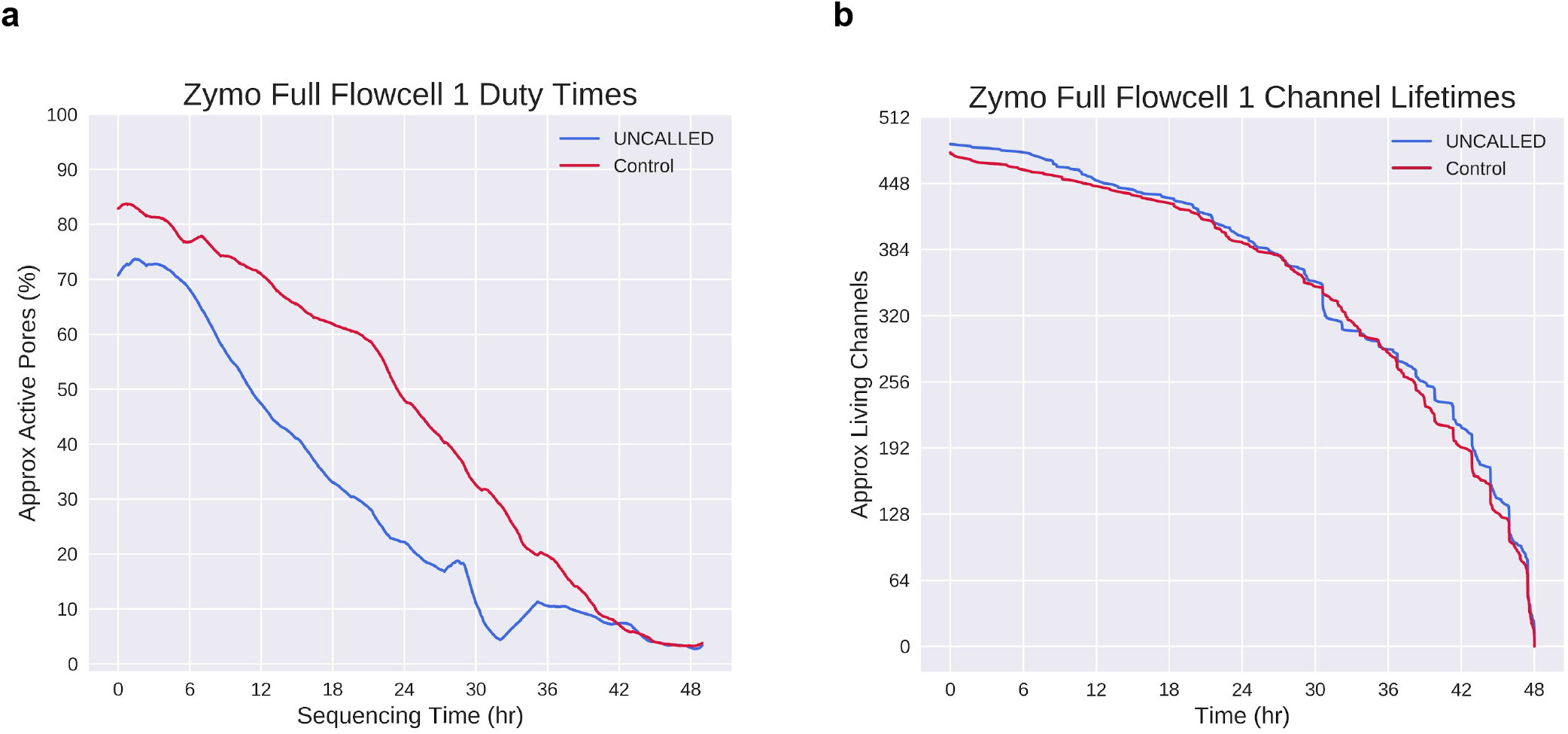
Pore activity during Zymo “full flowcell 1” sequencing runs. **(a)** Percent of channels that are labeled active throughout zymo bacterial depletion UNCALLED and control runs, based on the percent of signal labeled “pore” or “strand” in the MinKNOW duty times. Curves are smoothed by taking the mean of 92 minute windows, which smooths over mux scans. **(b)** Number of channels which are “alive” throughout the run, meaning they have the capacity to sequence reads, based on when the last read was produced. This is distinct from the duty time plots in that a channel may not produce a read for several hours but still be considered “alive”.

**Supplemental Figure S4.**
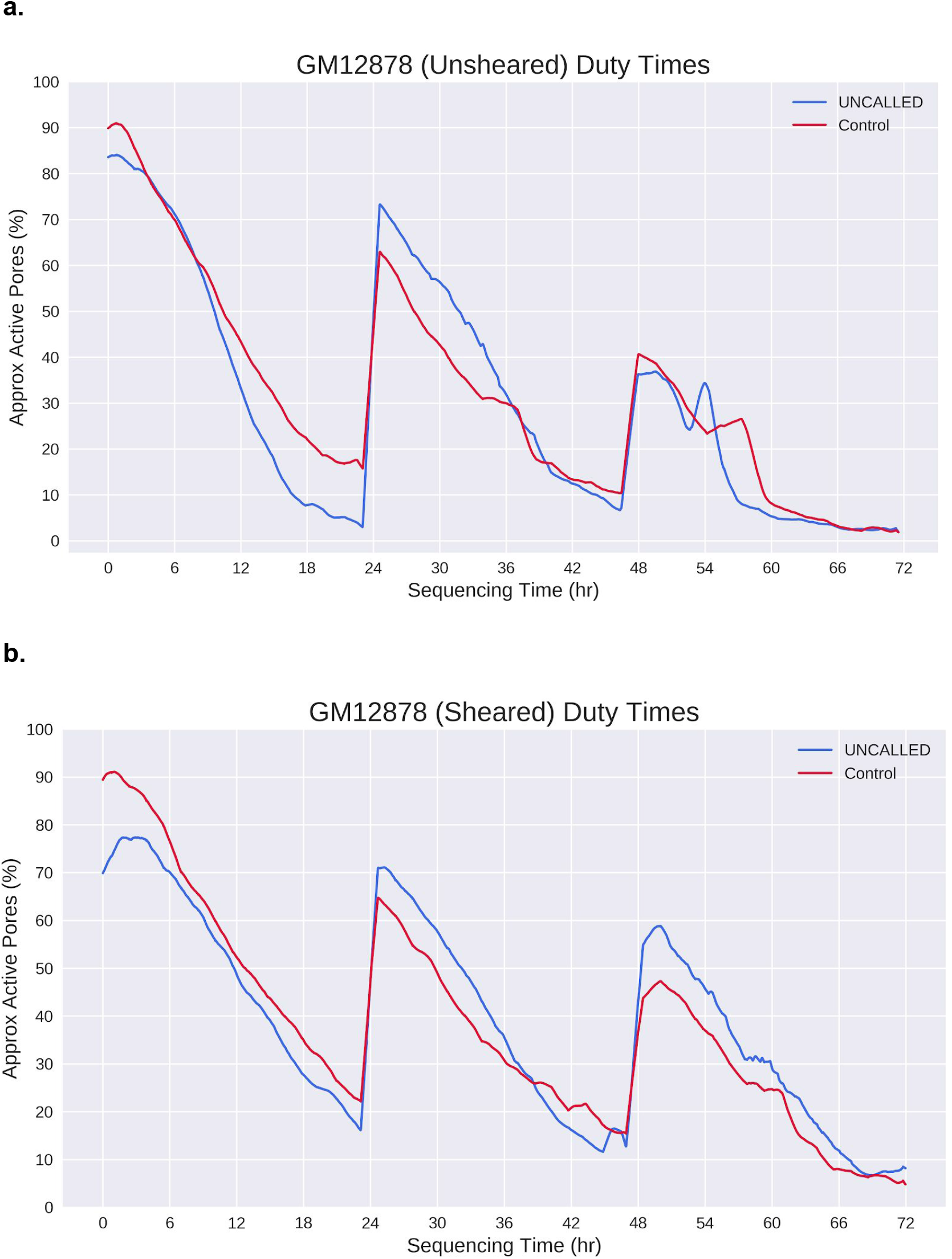
GM12878 gene enrichment run duty times in the **(a)** sheared run and **(b)** sheared run. Nuclease flushes were carried out at 24 and 48 hours in both runs. Curves plotted as in **Supplemental Fig. S3**. Note: we observed that a large patch of channels were marked as inactive after the second flush in the sheared UNCALLED run, which can occur because of bubbles introduced when loading.

**Supplemental Figure S5.**
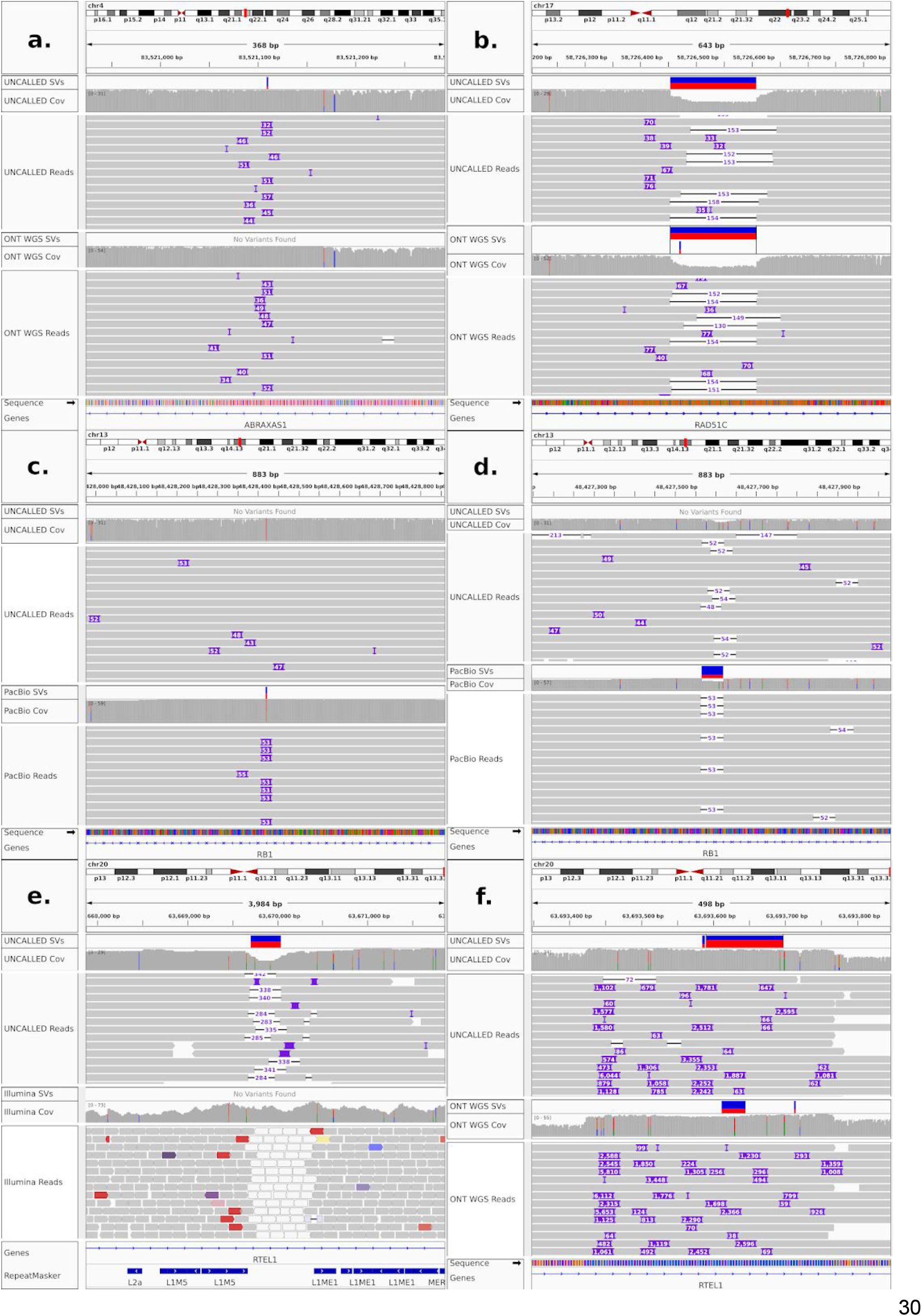
SVs confirmed by applying sensitive parameters in Sniffles and SURVIVOR or which required manual inspection to correct. **(a)** Insertion detected by UNCALLED but not by ONT WGS because most reads represented it as < 50bp. **(b)** Insertion detected by ONT WGS but not by UNCALLED because of low-complexity sequence. The overlapping deletion on the other haplotype also likely made the insertion difficult to resolve. **(c)** Insertions detected by UNCALLED but not by PacBio because of low-complexity sequence. **(d)** Deletion detected by PacBio but not by UNCALLED. **(e)** Deletion detected by UNCALLED (and all other long-read datasets) but not by Illumina reads, likely because of surrounding repetitive elements. Note that white read alignments indicate low mapping quality. **(f)** Sniffles called two SVs in this locus in both UNCALLED and ONT WGS, while it appears to represent a single duplication. SURVIVOR merged the ONT WGS SVs but not the UNCALLED SVs, causing a falsely unmatched SV. This is a known issue with SURVIVOR and this case was manually corrected.

**Supplemental Figure S6.**
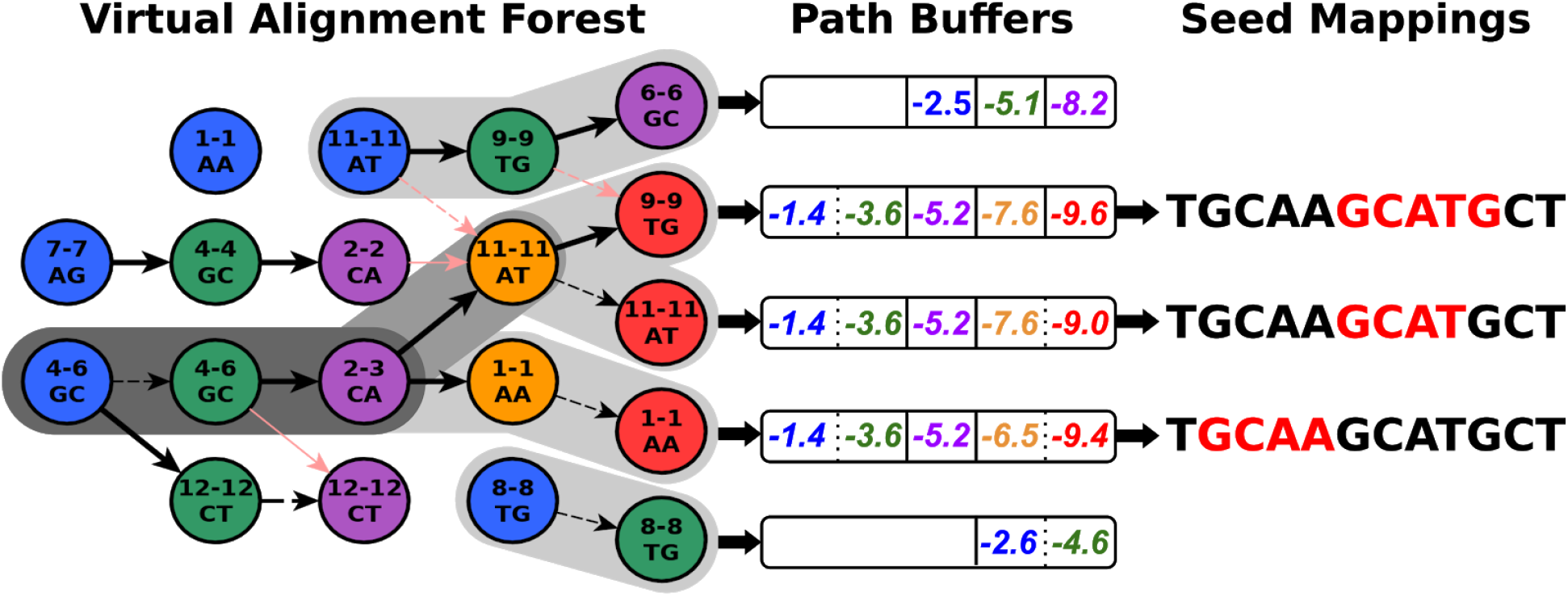
Representation of alignments in path buffers. The “Virtual Alignment Forest” is a more detailed version of the one in **Fig. 1a**. Pink edges mark paths that were pruned out due to lower probability in order to maintain the tree structure. Shaded backgrounds mark paths that have not been pruned out and are therefore represented in path buffers, and darker shading indicates that part of the path is represented in multiple buffers. “Path Buffers” store cumulative log probabilities that can be used to compute a rolling mean log probability as mapping progresses, as well as “stay” versus “move” events represented by dotted versus solid lines. Seed mappings are inferred from the FM index coordinate which are also stored in the buffers.

